# Genetic Diversity of *Hippophae rhamnoides* Varieties with Different Fruit Characteristics

**DOI:** 10.1101/2024.12.10.627738

**Authors:** Nataliya V. Melnikova, Alexander A. Arkhipov, Yury A. Zubarev, Roman O. Novakovskiy, Anastasia A. Turba, Elena N. Pushkova, Daiana A. Zhernova, Anna S. Mazina, Ekaterina M. Dvorianinova, Elizaveta A. Sigova, George S. Krasnov, Chengjiang Ruan, Elena V. Borkhert, Alexey A. Dmitriev

## Abstract

*Hippophae rhamnoides* is a valuable crop whose fruits are rich in bioactive compounds with health benefits. To date, there is a lack of genetic data for varieties of sea buckthorn. This fact hinders the identification of genetic determinants of valuable traits and limits the efficiency of breeding. In the present study, we analyzed a representative set of 55 valuable *H. rhamnoides* varieties of Russian breeding with different fruit characteristics and diverse lineages. Whole-genome sequencing was performed on the Illumina platform and at least 25× genome coverage was obtained for each accession. Based on the sequencing data, DNA polymorphisms were identified in genome regions corresponding to genes. These polymorphisms were used to evaluate the genetic relationships of the studied sea buckthorn varieties. We revealed genetically distinct groups of accessions that mostly corresponded to the lineages of the genotypes. Our data are important for assessing the effect of selection on sea buckthorn diversity and for evaluating the genetic relationship of different varieties, which is useful for breeders when selecting parental forms for crosses. The obtained information on DNA polymorphisms is also necessary to study the diversity of genes, including those that may determine valuable sea buckthorn traits, including fruit characteristics. Thus, our data can benefit both basic and applied research on sea buckthorn.

## 1 Introduction

Sea buckthorn (*Hippophae rhamnoides* L.) is a woody oil tree known for its fruits, which are a rich source of bioactive compounds, including carotenoids and flavonoids (Ciesarova et al., 2020; Mihal et al., 2023). In addition, the unique fatty acid composition of the fruit pulp oil, especially the high content of omega-7 monounsaturated palmitoleic acid, which is rare in plants, contributes to the nutritional benefits of its products (Sola Marsinach and Cuenca, 2019). In this regard, sea buckthorn products are used in medicine, cosmetics, and nutraceuticals (Gatlan and Gutt, 2021; Guo et al., 2022; Zuchowski, 2023). In addition to cultivation for fruit production, sea buckthorn is also used for ecological restoration due to its high resistance to extreme conditions (Ruan et al., 2013).

Sea buckthorn is mainly cultivated in China (2.07 million ha), India (0.02 million ha), Romania (0.02 million ha), Mongolia (0.02 million ha), Russia (0.01 million ha), and Pakistan (0.01 million ha) (Nybom et al., 2023). Thus, 90% of sea buckthorn resources are located in China (Singh, 2022). However, the pioneer in sea buckthorn breeding was Russia, where selection of *H. rhamnoides* ssp. *mongolica* Rousi started in 1933 and allowed the development of a wide range of high-yield varieties with high-quality fruits (Singh and Zubarev, 2014). In contrast, breeding of sea buckthorn in China started later, mainly with *H. rhamnoides* ssp. *sinensis* Rousi (Nybom et al., 2023). Varieties of *H. rhamnoides* ssp. *mongolica* are characterized by large fruits and high yield, high oil content, and lower acidity compared to *H. rhamnoides* ssp. *sinensis* varieties, which are better adapted to abiotic and biotic stressors (Nybom et al., 2023). Sea buckthorn breeding does not stand still, new improved varieties are being developed. The use of genetic data can improve the efficiency of sea buckthorn breeding.

To date, high-quality genome assemblies of *H. rhamnoides* with sizes of 849, 730, and 919 Mb were obtained (Wu et al., 2022; Yu et al., 2022; Yang et al., 2024). In addition, the genomes of *Hippophae tibetana* (957 and 1453 Mb) (Wang et al., 2022b; Zhang et al., 2024) and *Hippophae gyantsensis* (716 Mb) (Chen et al., 2024) were assembled. Moreover, whole-genome sequencing of 40 wild *H. rhamnoides* ssp. *mongolica* and *H. rhamnoides* ssp. *sinensis* representatives and 15 cultivated *H. rhamnoides* ssp. *mongolica* varieties was performed in China (Yu et al., 2022). Therefore, there is a lack of genomic data for varieties of sea buckthorn. The aim of the present study was to fill this gap by performing whole-genome sequencing of the unique set of 55 varieties of Russian breeding, which are likely to be significantly different from the Chinese varieties.

## 2 Materials and Methods

### 2.1 Plant Material

A set of 56 accessions of *Hippophae rhamnoides* L. was formed to cover the diversity of sea buckthorn cultivated in Russia: one replicate for 54 varieties and two biological replicates for Elizaveta. The following valuable characteristics were considered: weight, flavor, shape, and color of fruits and differences in origin (Table 1). Characteristics of sea buckthorn varieties were assessed according to Kondrashov et al. (Kondrashov et al., 1999). Dormant shoots of the selected genotypes were collected at the Federal Altai Scientific Center of Agrobiotechnologies (Barnaul, Russia) in April 2023. The shoots were placed in containers with water in a room with a temperature of ∼22 °C. After the leaves appeared, they were collected in tubes, frozen in liquid nitrogen, and further stored in a low-temperature freezer until DNA extraction.

**Table 1.**
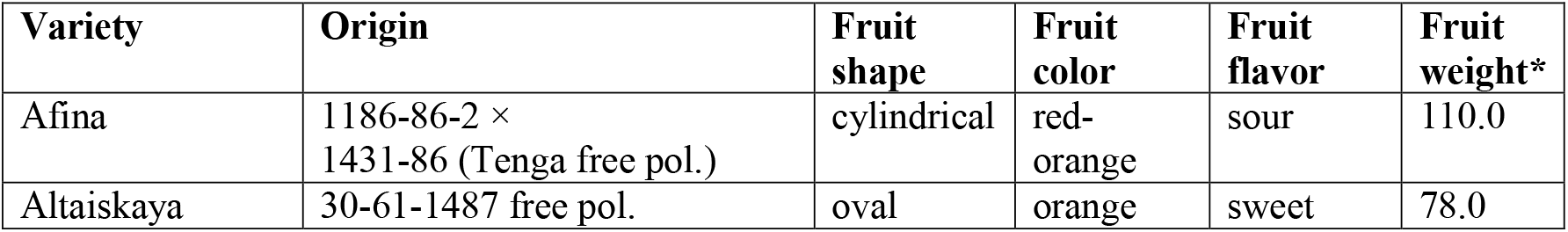
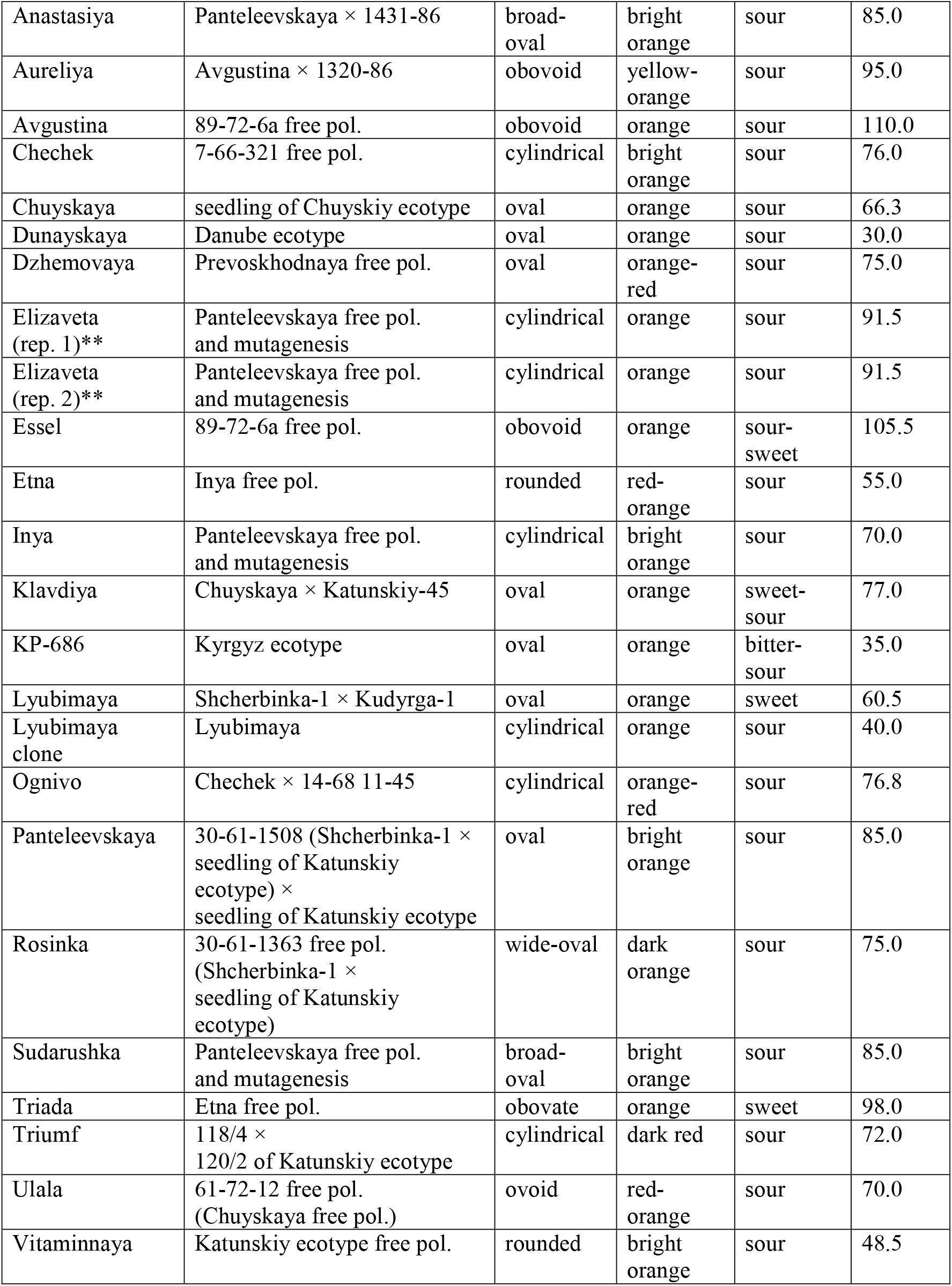
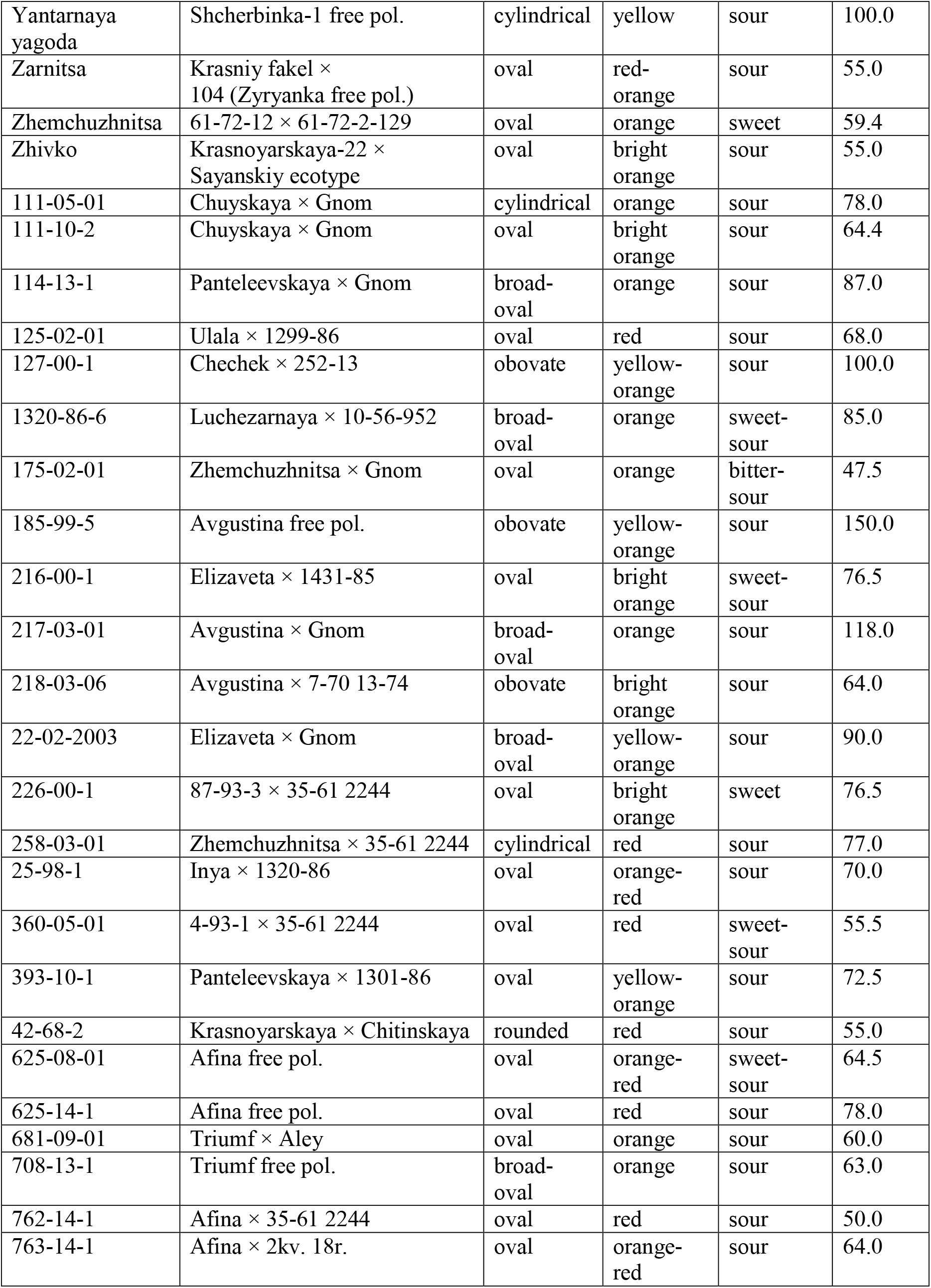
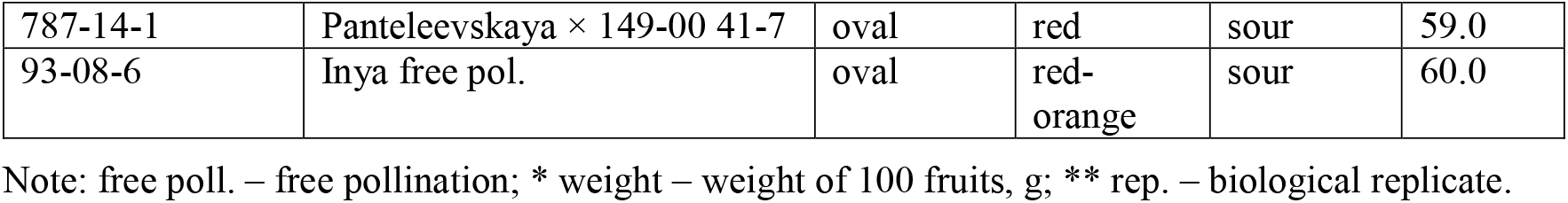
Characteristics of sea buckthorn varieties analyzed in the study.

### 2.2 DNA Extraction

DNA was extracted using the Magen HiPure Plant DNA Mini Kit (Magen, Guangzhou, China). The quality and quantity of DNA were evaluated using NanoDrop 2000C (Thermo Fisher Scientific, Waltham, MA, USA), Qubit 4.0 (Thermo Fisher Scientific), and agarose gel electrophoresis (2% agarose).

### 2.3 Whole-Genome Sequencing

The QIAseq FX DNA Library UDI Kit (Qiagen, Chatsworth, CA, USA) was used for DNA library preparation. Quantity and quality of DNA libraries were assessed using Qubit 4.0 (Thermo Fisher Scientific) and Qsep1-Plus (Bi-Optic, New Taipei City, Taiwan). Genome sequencing was performed on a NovaSeq 6000 (Illumina, San Diego, CA, USA) with a read length of 150 + 150 bp.

### 2.4 Sequencing Data Analysis

The obtained Illumina reads were processed with Trimmomatic 0.39 (TRAILING:28, SLIDINGWINDOW:4:17, MINLEN:40) (Bolger et al., 2014). The processed reads were mapped to the annotated *H. rhamnoides* genome from the CNGB Nucleotide Sequence Archive (https://db.cngb.org/cnsa), Project ID CNP0001846 (Wu et al., 2022), and VAF (Variant Allele Frequencies) values were calculated for genome regions corresponding to genes (exons and introns) using PPLine (Krasnov et al., 2015). Genetic distances between sea buckthorn varieties were calculated and clustered with Ward’s method (ward.D2) in PPLine (Krasnov et al., 2015).

## 3 Preliminary Data Analysis

Representative set of 55 sea buckthorn varieties was formed from the unique collection of the Federal Altai Scientific Center of Agrobiotechnologies (Barnaul, Russia). The selected varieties had different fruit characteristics and different origins in order to maximize the diversity of the analyzed set (Table 1).

Whole-genome sequencing was performed and at least 23 Gb of raw Illumina data (150 + 150 bp reads) were obtained for each accession, which corresponded to more than 25× genome coverage (raw Illumina reads were deposited to NCBI SRA, BioProject PRJNA1177110). After mapping the reads to the annotated *H. rhamnoides* reference genome, data on about 4 million DNA polymorphisms in genes were obtained (lists of DNA polymorphisms were deposited to Zenodo, https://zenodo.org/records/13999625). These data are useful for studying the diversity of allelic variants for specific genes, especially those that may be associated with valuable traits, such as the content of bioactive compounds and other fruit characteristics and resistance to stressors. It is worth noting that a significant part of the identified DNA polymorphisms was present in all analyzed sea buckthorn varieties, indicating that they are genetically distinct from the used reference genome. In addition, genetic distances between the accessions were calculated to evaluate their relationships (Supplementary Table 1).

To visualize the relationships of the studied varieties based on DNA polymorphisms in gene sequences, a dendrogram was constructed (Figure 1). Cluster I was the most distinct and included KP-686 (Kyrgyz ecotype), Dunayskaya, and Yantarnaya Yagoda, which are not varieties of Altai breeding and probably have significant differences at the genome level from the other studied accessions. The same cluster included 175-02-01, obtained by crossing varieties of Altai breeding, and its position in the dendrogram is not expected and requires additional research.

**Figure 1.**
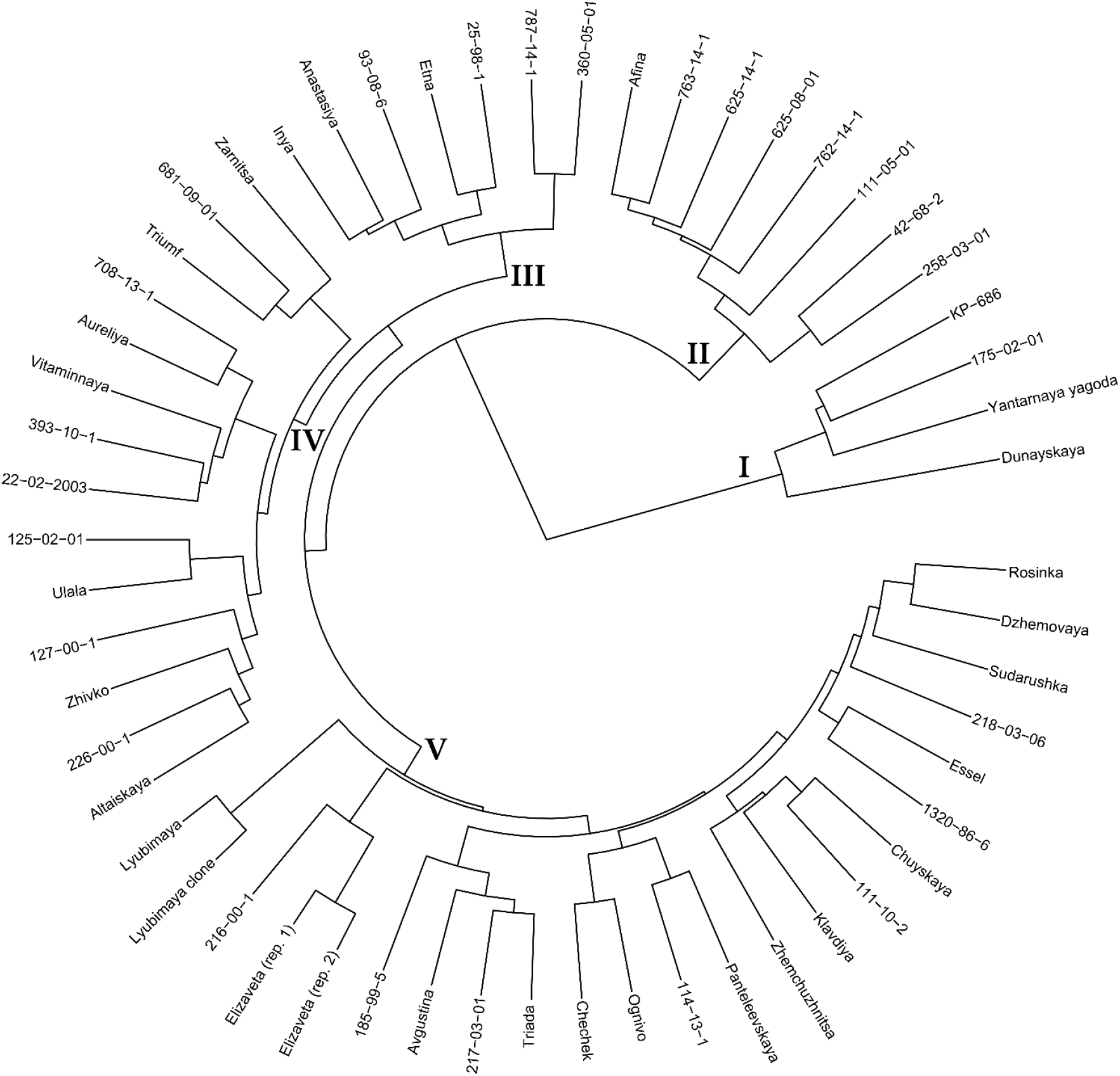
Clustering of sea buckthorn varieties based on the data of whole-genome sequencing. DNA polymorphisms (VAF values) in gene sequences were analyzed. Ward’s method of cluster analysis.

The remaining sea buckthorn varieties were divided into four clusters. Cluster II included Afina and all studied progenies of this variety, namely 625-08-01, 625-14-1, 762-14-1, and 763-14-1. It can be assumed that Afina and sea buckthorn genotypes obtained with its participation are genetically quite different from the other studied varieties of Altai breeding. In addition, 111-05-01, 258-03-01, and 42-68-2, which are believed to be unrelated to Afina, were in Cluster II, which is difficult to explain from a genealogical point of view.

Cluster III clearly distinguished a group of sea buckthorn varieties with Panteleevskaya in their lineages. Thus, this group is likely to be significantly different from other studied sea buckthorn genotypes at the genome level.

Cluster IV included 14 varieties, among which the genetic relationships were not as clear as in the first three clusters, but they were still present. So, a group of four Novosibirsk accessions was isolated: Triumf, Zarnitsa, 681-09-01, and 708-13-1, with Triumf being the parental form for 681-09-01 and 708-13-1. Ulala and its progeny 125-02-01 were also in this cluster. Two varieties with Panteleevskaya in their lineages were also in Cluster IV: 22-02-2003 and 226-00-01. In general, however, this cluster contained a mixture of quite different sea buckthorn varieties.

Cluster V contained 23 accessions. In this cluster, as in other clusters, some relationships corresponding to lineages were observed. For example, Rosinka and Sudarushka, which entered this cluster, have common roots. In addition, varieties Essel and 218-03-06 have the genotype 89-72-6a in their lineages. 89-72-6a is very interesting in terms of strong inheritance of large fruit size. In this respect, it is the progenitor of many varieties, most of which were present in Cluster V. The exception was the variety Aureliya, which was in Cluster IV. Other relationships can also be traced in Cluster V. For example, Lyubimaya clone is a seedless mutant of the variety Lyubimaya. Several closely related groups were also isolated: Elizaveta (two biological replicates) and its progeny 2016-00-1, Chechek and its progeny Ognivo, Chuyskaya and its progenies Klavdiya and 111-10-2, and Panteleevskaya and its progeny 114-13-1.

In general, the dendrogram we obtained based on DNA polymorphisms in all sea buckthorn genes annotated in the used reference genome (Wu et al., 2022) reflected well the known data on the relationship of the studied genotypes. The research on *H. rhamnoides* performed by Yu et al. using whole-genome sequencing allowed them to separate wild genotypes of *H. rhamnoides* ssp. *mongolica* from cultivated ones, as well as to separate *H. rhamnoides* ssp. *sinensis* accessions into a separate group (Yu et al., 2022). However, we were unable to find any other work that characterized representative sets of sea buckthorn genotypes using whole-genome sequencing (NCBI PubMed, https://pubmed.ncbi.nlm.nih.gov/; Google Scholar, https://scholar.google.com; accessed October 28, 2024). Meanwhile, whole-genome sequencing and linkage mapping is an urgent need for sea buckthorn studies (Sharma, 2022).

Data on the diversity of sea buckthorn varieties at the genome level are of great value for understanding the extent to which selection affected the gene pool of this crop, and what patterns can be traced by analyzing the genetic data. We studied the sea buckthorn varieties of Russian breeding, which has a long history. The forms with valuable traits created by Russian breeders became the progenitors of many varieties all over the world (Singh, 2022), so the obtained by us data are of special value. In addition, the evaluation of genetic relationships of different accessions is important for breeders when selecting parental forms for crosses.

Recently, there has been an increasing number of articles devoted to the beneficial properties of sea buckthorn (Wang et al., 2022a; Chen et al., 2023; Mihal et al., 2023; Nybom et al., 2023; Teng et al., 2024; Xu et al., 2024), but in terms of genetics, this crop is still relatively understudied (Sharma, 2022). Indeed, several high-quality genome assemblies of *H. rhamnoides* were obtained (Wu et al., 2022; Yu et al., 2022; Yang et al., 2024) and some transcriptome studies were performed (Bansal et al., 2018; Ye et al., 2018; Gao et al., 2022; Lyu et al., 2022; Yu et al., 2022). A number of works were also devoted to fatty acid synthesis in sea buckthorn and genes/microRNAs involved in this process (Ding et al., 2018; Ding et al., 2019; Ding et al., 2022; Yu et al., 2022). However, the genetic determinants and their diversity remain unknown for most of the key traits that define the value of sea buckthorn varieties, including carotenoid content, fruit shape and flavor. In this context, data on DNA polymorphisms in gene sequences obtained for a representative set of accessions characterized by phenotype will allow the search for associations between allelic variants of genes and valuable traits. These data are the basis for the development of marker-assisted and genomic selection of sea buckthorn, which are increasingly used in breeding practice for other agricultural plants (Xu et al., 2020; Hasan et al., 2021; Thudi et al., 2021; Dmitriev et al., 2022; Werner et al., 2023; Mangal et al., 2024).

## 5 Author Contributions

N.V.M., Y.A.Z., and A.A.D. conceived and designed the work. Y.A.Z., R.O.N., A.A.T., E.N.P., D.A.Z., A.S.M., and E.V.B. performed the experiments. N.V.M., A.A.A., Y.A.Z., E.M.D., E.A.S., G.S.K., C.R., and A.A.D. analyzed the data. N.V.M., Y.A.Z., and A.A.D. wrote the manuscript. All authors read and approved the final manuscript.

## 7 Acknowledgments

We thank the Center for Precision Genome Editing and Genetic Technologies for Biomedicine, EIMB RAS for providing computing power and techniques for the data analysis. This work was performed using the equipment of the EIMB RAS “Genome” center (http://www.eimb.ru/ru1/ckp/ccu_genome_ce.php).

## 8 Supplementary Material

**Supplementary Table 1**. Matrices of genetic distances between 56 sea buckthorn accessions based on the analysis of DNA polymorphisms in gene sequences at the whole-genome level.

## 9 Data Availability Statement

The datasets generated for this study can be found in the NCBI SRA database under the BioProject accession number PRJNA1177110.

